# Single-cell transcriptomics uncovers chromatin dysfunction in a human TDP-43 proteinopathy model of Amyotrophic Lateral Sclerosis

**DOI:** 10.64898/2026.05.10.724071

**Authors:** Johanna Ganssauge, Harshitha B Chandrashekar, Yiyun Zhang, Antonio R Fusciardi, Seema C Namboori, Akshay Bhinge

## Abstract

TDP-43 proteinopathy, characterised by nuclear depletion and cytoplasmic aggregation of TDP-43, is the defining pathological hallmark of amyotrophic lateral sclerosis (ALS) and a shared pathology across frontotemporal lobar degeneration with TDP-43 inclusions (FTLD-TDP), limbic-predominant age-related TDP-43 encephalopathy (LATE), and a substantial subset of Alzheimer’s disease. We recently developed a human model of TDP-43 proteinopathy that enables inducible mislocalisation of endogenous TDP-43 in iPSC-derived neurons without chemical stress or mutant protein overexpression.

Using single-cell RNA sequencing of this model, we dissected the transcriptomic consequences of TDP-43 nuclear depletion across motor neurons as well as V1 and V2 interneurons at single-cell resolution. This approach uncovered disruption of ATP-dependent chromatin remodelling as a convergent downstream pathway across all three spinal neuron subtypes. Master regulator analysis identified ACTL6B, the neuron-specific subunit of the nBAF (neuronal BRG1/BRM-associated factor) chromatin remodelling complex, as the most consistently inhibited transcription factor following TDP-43 mislocalisation. ACTL6B downregulation emerges early in the mislocalisation cascade and is confirmed in post-mortem ALS spinal cord. *ACTL6B* knockdown in post-mitotic motor neurons phenocopies both the morphological and transcriptional consequences of TDP-43 pathology. Together, these findings establish nBAF complex dysfunction as a principal, spinal cord-enriched driver of TDP-43-associated neurodegeneration and reveal chromatin remodelling defects as a key mechanism in ALS.

## Introduction

ALS is a fatal neurodegenerative disease characterised by the selective loss of upper and lower motor neurons, leading to progressive paralysis and death from respiratory failure, typically within three to five years of diagnosis (Brown and Al-Chalabi, 2017). There are no effective treatments. ALS is classified as sporadic (sALS, ∼85% of cases) or familial (fALS, ∼15%), with mutations in over 30 genes linked to fALS, including *C9ORF72*, *SOD1*, *TARDBP*, and *FUS* (Brown and Al-Chalabi, 2017; Hardiman et al., 2017). The mechanistic boundaries between sALS and fALS are increasingly blurred; TDP-43 proteinopathy, pathological nuclear clearance and cytoplasmic aggregation of TDP-43, is a neuropathological hallmark shared across the vast majority of ALS cases regardless of aetiology (Neumann et al., 2006).

TDP-43 is a predominantly nuclear RNA-binding protein that regulates pre-mRNA splicing, mRNA stability, and transport (Ling et al., 2013). Its nuclear depletion in ALS leads to widespread transcriptomic dysregulation, most strikingly through the derepression of cryptic exon splicing. Under normal conditions, TDP-43 suppresses the inclusion of unannotated intronic exons; when depleted, these cryptic exons are incorporated into mature transcripts, frequently disrupting protein function (Ling et al., 2015). Cryptic exon inclusion in *STMN2* and *UNC13A* are now well-validated molecular signatures of TDP-43 loss-of-function in ALS patient neurons (Klim et al., 2019; Melamed et al., 2019; Ma et al., 2022).

While bulk RNA sequencing has illuminated global transcriptomic shifts in ALS, its utility in dissecting cell-type-specific transcriptomic alterations is inherently limited by cellular heterogeneity (Namboori et al., 2021). Conversely, single-cell RNA sequencing offers a high-resolution approach to investigate transcriptomic changes in specific cell populations affected by TDP-43 pathology, particularly in vulnerable neuronal subtypes (Gittings et al., 2023). This cell-type-specific resolution is crucial for identifying the precise chromatin remodellers and associated regulatory pathways impacted by TDP-43 dysfunction, which are often masked in bulk tissue analyses (Li et al., 2023; Rizzuti et al., 2023). For example, single-nucleus sequencing has been used to analyse neuronal nuclei with and without TDP-43 inclusions from post-mortem human brain tissue, allowing characterisation of cryptic exon inclusion events in specific cell types (Gittings et al., 2023). This approach has revealed that specific neuronal populations, such as excitatory cortical neurons, are particularly affected by TDP-43 pathology, exhibiting distinct transcriptional aberrations including cell-type-specific cryptic exon inclusion events and disease-associated epigenetic alterations (Ruf et al., 2026).

Chromatin remodelling dysfunction is an emerging but underappreciated driver of neurodegeneration in ALS. The neuronal BAF (nBAF, neuronal BRG1/BRM-associated factor) complex, polycomb repressive complexes, and related chromatin remodellers maintain the transcriptional programmes essential for neuronal identity, axonal integrity, and stress responses in post-mitotic neurons (Wu et al., 2007, Schimmelmann et al., 2016). While aberrant histone modifications and chromatin remodelling dysregulation have been reported in ALS patient tissue (Tibshirani et al., 2017), whether TDP-43 pathology directly compromises chromatin remodelling has not been systematically addressed.

Here, we use single-cell RNA sequencing of iPSC-derived spinal neurons in which endogenous TDP-43 mislocalisation is induced without chemical stress or mutant protein overexpression (Ganssauge et al., 2025) to establish the transcriptomic consequences of TDP-43 proteinopathy at cell-type resolution.

## Results

### Differentiation of iPSC-derived motor neurons and induction of TDP-43 mislocalisation

We deployed our recently developed human model of TDP-43 proteinopathy in which endogenous GFP-tagged TDP-43 is inducibly mislocalised in iPSC-MNs through doxycycline-driven expression of an anti-GFP nanobody fused to a nuclear export signal (NES). In parallel MN cultures, expression of the nanobody without the NES serves as control resulting in nuclear-retained TDP-43 (Control). This results in controlled cytoplasmic accumulation of TDP-43 in iPSC-derived MNs, leading to dendritic defects, increased apoptosis and splicing changes that correlate with the degree of cytoplasmic TDP-43 accumulation (Ganssauge et al., 2025). To induce TDP-43 mislocalisation, doxycycline was added to the culture medium at day 20 of differentiation to express the anti-GFP nanobodies and cells were subsequently harvested at two time points, day 22 and day 40 (Fig.1A), representing an early and a more extended period of TDP-43 cytoplasmic mislocalisation, respectively, allowing us to capture both the immediate transcriptomic response and the consequences of sustained proteinopathy within the same experimental framework.

**Figure 1.**
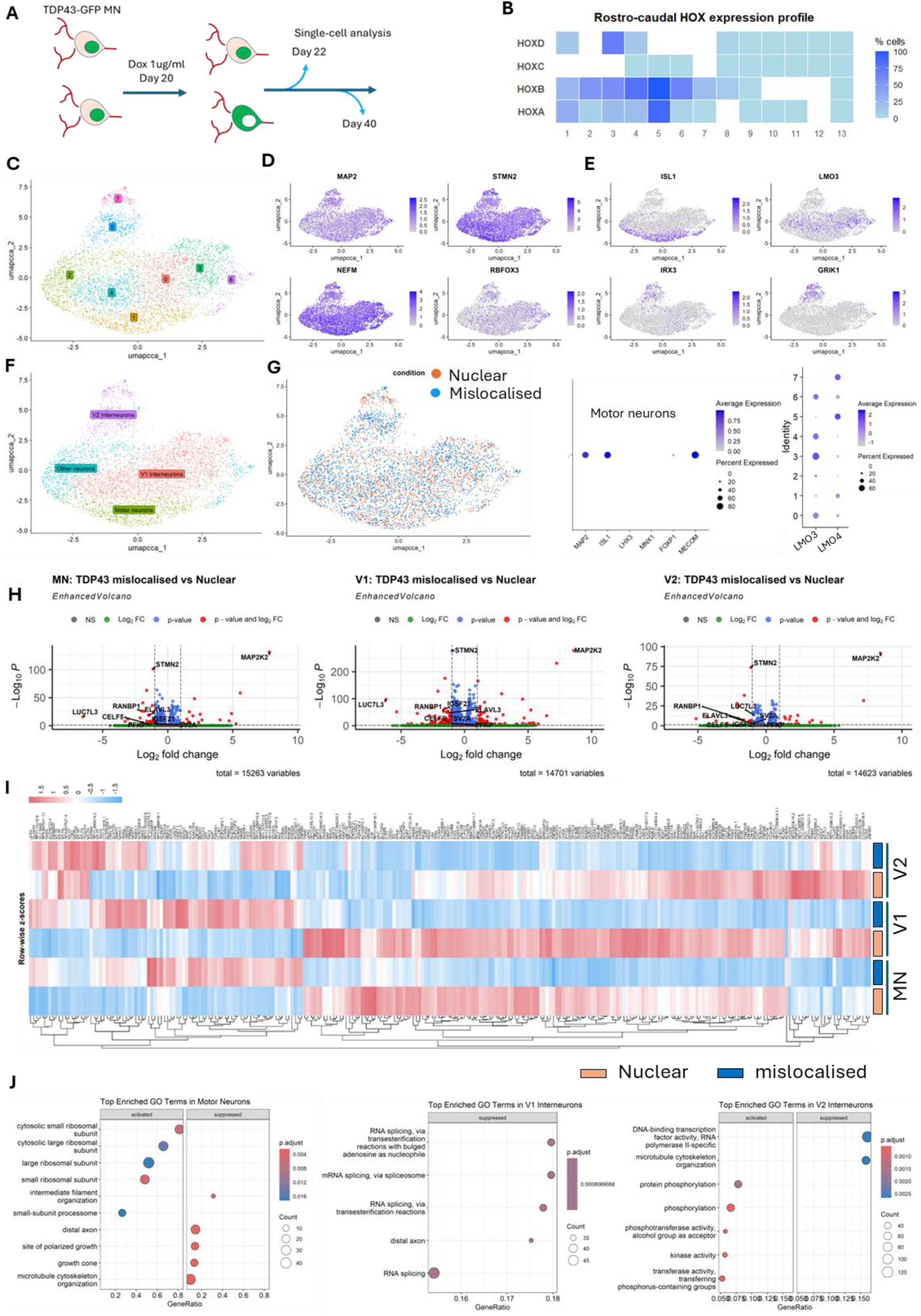
Single-cell transcriptomic profiling of iPSC-derived spinal neurons following TDP-43 mislocalisation. (A) Schematic of the experimental design. TDP-43-GFP iPSCs were differentiated into spinal neurons and doxycycline was added at day 20 to induce TDP-43 mislocalisation. Cells were harvested at day 22 and day 40 for single-cell RNA sequencing using PIP-seq. (B) Heatmap showing the percentage of cells expressing each HOX gene across paralog groups 1–13. The majority of cells express HOXA5 and HOXB5, with sparse expression of HOXB8 and HOXD8 and no expression of paralog group 9 or higher, indicating a hindbrain to brachial spinal cord identity. (C) UMAP of the integrated single-cell dataset showing distinct transcriptional clusters after CCA intergration. (D) Feature plots showing normalised expression of pan-neuronal marker genes across the UMAP. (E) Feature plots showing expression of subtype-specific markers used to annotate clusters as motor neurons (MNs) [ISL1], V1 interneurons [LMO3], and V2 interneurons [IRX3 and GRIK1]. (F) UMAP with cluster annotations. Cells that could not be unambiguously assigned to a known spinal neuronal subtype are designated as other neurons. (G) UMAP coloured by condition, showing that control (nuclear TDP-43) and mislocalised cells are evenly distributed across all clusters, confirming that clustering is driven by cell identity rather than condition. (H) Volcano plots showing differential gene expression in MNs, V1 interneurons, and V2 interneurons comparing TDP-43 mislocalised versus control cells. Genes meeting both fold change (absolute fold change > 2) and adjusted p-value thresholds < 0.01 are highlighted in red. ALS-associated genes are labelled. Total number of genes tested is indicated for each cell type. (I) Heatmap showing row-wise z-scored normalised expression of differentially expressed genes across MN, V1, and V2 interneuron subtypes. Samples are grouped by condition and cell type. Colour bar indicates subtype identity. (J) Dot plots showing the top enriched Gene Ontology biological process terms in the activated and suppressed DEG sets for MNs, V1 interneurons, and V2 interneurons. Dot size indicates gene count and colour indicates adjusted p-value.

Single-cell RNA sequencing libraries were generated using PIP-seq (Fluent Biosciences). A total of 5139 cells were captured at day 22 and 3939 cells at day 40 across all conditions and differentiations prior to quality filtering.

### Quality control and filtering of single-cell transcriptomic data

Raw sequencing data were processed through a rigorous quality control pipeline to ensure downstream analyses were performed on high-confidence, cell-associated transcriptomes. Ambient RNA contamination, a pervasive artefact in droplet-based single-cell sequencing arising from RNA released by lysed cells, was first estimated and removed using DecontX, a Bayesian decontamination method (Yang et al., 2020). DecontX was applied to the filtered barcode-cell matrix using the raw, unfiltered matrix as a background reference. To maximise the accuracy of ambient RNA estimation and minimise the risk of overcorrection, the background matrix was restricted to barcodes with fewer than 100 unique molecular identifiers (UMIs), thereby enriching for true empty droplets rather than low-quality cells.

Following ambient RNA correction, low-quality cells were identified and excluded based on three standard metrics: the total number of UMIs detected per cell, the total number of genes detected, and the percentage of reads mapping to the mitochondrial genome. Elevated mitochondrial read fractions are a well-established indicator of cellular stress or membrane compromise, and cells failing thresholds on any of these metrics were removed from further analysis (Ilicic et al., 2016; Luecken and Theis, 2019). Additionally, doublet detection was performed computationally (McGinnis et al., 2019); on average, fewer than 3% of barcodes were identified as doublets, consistent with expected rates for the cell input used, and these were excluded prior to downstream processing. After applying all quality filters and removing doublets, we retained a total of 4741 high-quality single-cell transcriptomes at day 22 and 3289 at day 40.

### HOX gene expression profiling reveals a hindbrain and brachial spinal cord identity

A fundamental consideration when interpreting iPSC-derived MN data in the context of spinal cord biology is the rostro-caudal identity of the neurons produced in vitro. In the developing spinal cord, MNs at distinct rostro-caudal levels are specified by combinatorial expression of HOX transcription factors, termed the HOX code, which determines the connectivity, target musculature, and disease vulnerability of different MN pools (Dasen et al., 2005; Dasen and Jessell, 2009). To characterise the rostro-caudal address of our in vitro-differentiated neurons, we systematically estimated the proportion of cells expressing each of the 39 HOX genes across paralog groups 1–13.

This analysis revealed that the majority of cells expressed HOXA5 and HOXB5, markers associated with the hindbrain and brachial (cervical) spinal cord, while expression of HOXB8 and HOXD8, characteristic of more caudal, thoracic levels, was detected in only a minor fraction of cells (Fig.1B). No cells expressed HOX genes from paralog groups 9 and above, which mark lumbar and sacral spinal cord identities (Dasen and Jessell, 2009). Taken together, this HOX expression profile indicates that our differentiation protocol consistently generates neurons with a rostral, hindbrain-to-brachial spinal cord identity, broadly consistent with the MN populations that innervate the upper limb musculature and are amongst the earliest and most severely affected in ALS.

### Unsupervised clustering resolves distinct neural subtypes in iPSC-derived spinal cultures

Initial unsupervised clustering of the ambient RNA-corrected, quality-filtered single-cell matrix revealed that cells tended to cluster by batch and sample of origin rather than by biologically meaningful transcriptional state, indicating the presence of technical variation that would confound downstream interpretation. To address this, we systematically benchmarked multiple normalisation and integration strategies, including log-normalisation, SCTransform (Hafemeister and Satija, 2019), canonical correlation analysis (CCA) integration (Stuart et al., 2019), and Harmony (Korsunsky et al., 2019). Evaluation of each approach based on its ability to dissolve batch structure whilst preserving biological signal led us to select log-normalisation followed by CCA integration as the optimal combination that most effectively aligned cells across batches without over-correcting genuine transcriptional heterogeneity.

For the day 22 dataset, the two independent differentiations were first merged using CCA integration prior to clustering. We then performed iterative clustering by incrementally increasing the Leiden resolution parameter from 0.6 to 1.2 in steps of 0.2, assessing the biological coherence of the resulting clusters at each step using a panel of established motor neuron (MN) and interneuron (IN) marker genes. At a resolution of 0.8, this approach yielded nine clusters, with the ninth cluster containing only 20 cells excluded from further analyses. The remaining eight clusters all expressed pan-neuronal markers *MAP2, STMN2*, *NEFM* and *RBFOX3* indicating that all cells were neurons (Fig.1C,D). Six of these clusters could be assigned to recognised spinal neuronal subtypes MNs (ISL1), V1 interneurons (LMO3) and V2 interneurons (GRIK1, IRX3) based on their marker expression profiles (Fig.1E, F). Increasing the resolution beyond 0.8 resulted in fragmentation of the MN clusters into sub-clusters that were widely dispersed in UMAP space without a corresponding gain in biological interpretability, suggesting over-partitioning of a transcriptionally coherent population. Resolution 0.8 was therefore selected for the final clustering of day 22 samples. Two of the eight clusters exhibited mixed expression of both MN and IN markers, including ISL1 and LMO3; these ambiguous clusters were termed “other neurons” and excluded from further analyses (Fig.1F). The remaining six clusters were annotated as MNs, V1 interneurons, and V2 interneurons based on their marker gene signatures (Fig.1F). Additionally, the majority of the MNs expressed MECOM, a marker of expressed in MMC MNs (Fig.1E). Similarly, expression of LMO4 and GRIK1 in the majority of the V2 IN population indicated these were V2b inhibitory interneurons (Fig.1E). Post-integration, cells with nuclear or mislocalised TDP-43 were evenly distributed across each neuronal subtype (Fig. 1G).

We applied a similar approach to classify the day 40 single-cell dataset. Given the increased transcriptional complexity expected at this later time point, reflecting more mature neuronal states and the compounding effects of sustained TDP-43 mislocalisation, optimal cluster resolution was achieved at 1.2, yielding 12 clusters in total. Replicate 1 of the day 40 dataset was found to contain significantly fewer cells post-filtering than replicate 2, indicative of a lower quality sample preparation; this replicate was therefore excluded and day 40 cells were re-clustered, yielding six clusters. As at day 22, clusters were annotated using known marker genes, and the same neuronal subtypes MN, V1, and V2 IN were successfully identified.

### TDP-43 mislocalisation drives widespread transcriptional dysregulation in neuronal subtypes

To detect early changes, we focused our analyses on the day 22 dataset. We performed cell-type-specific differential expression analysis using both single-cell Wilcoxon testing (fold change > 2, adjusted p < 0.05) and pseudobulk averaging to guard against spurious p-values arising from the non-independence of cells derived from the same sample (Crowell et al., 2020). V1 INs displayed the strongest response by both methods; 233 genes by Wilcoxon and 22 by pseudobulk, followed by MNs (71 and 10 genes, respectively), with V2 INs showing the most limited response (40 and 4 genes). This indicates that in addition to MNs, V1 INs are also sensitive to TDP-43 pathology whilst V2 INs are relatively resistant.

In MNs, *STMN2* was the most significantly downregulated gene (Figure 1H). Loss of STMN2 is a well established molecular signature of TDP-43 nuclear depletion in ALS patient tissue (Klim et al., 2019; Melamed et al., 2019), and its recovery here confirms our model faithfully recapitulates TDP-43 loss-of-function. Other downregulated genes included *ELAVL3, CELF5, RANBP1, IGSF21*, and *PFKP*, all previously linked to ALS, alongside *SV2A*, a synaptic vesicle glycoprotein reduced in ALS patients, and *LUC7L3*, a pre-mRNA splicing factor indicating disruption of core spliceosome machinery. *MAP2K2* was amongst the most significantly upregulated genes.

The V1 IN response closely mirrored that of MNs (Fig.1H). *STMN2* was again the most significantly downregulated gene, *LUC7L3* was similarly downregulated, and MAP2K2 was again amongst the most significantly upregulated, recapitulating the top MN hits with notable consistency. In V2 INs, *STMN2, ELAVL3*, and *RANBP1* were significantly downregulated and MAP2K2 upregulated, whilst *CELF5, IGSF21, LUC7L3*, and *PFKP* did not reach significance (Fig.1H), consistent with the more muted transcriptional response in this subtype (Fig.1I).

To enable threshold-independent pathway analysis, we performed GSEA across all three cell types. In MNs, downregulated pathways spanned multiple functional categories (Fig.1J): neuronal structure terms including distal axon and growth cone, consistent with the axonal degeneration characteristic of ALS, alongside RNA splicing and spliceosome terms indicating disruption of co-transcriptional RNA processing, protein folding, and chromatin organisation. Upregulated pathways in MNs were dominated by cytosolic ribosomal subunit genes. Rather than reflecting increased translational capacity, this likely represents a compensatory response to UPR activation: ER stress drives ribosomal biogenesis through ATF4-mediated transcription and transient mTORC1 activation as part of a cellular attempt to restore proteostatic balance (Pakos-Zebrucka et al., 2016). In V1 INs, downregulated pathways were similarly enriched for RNA splicing and axonal structure terms, with no significantly upregulated pathways identified (Fig.1J). V2 INs showed downregulation of axonal cytoskeletal and transcription factor-related terms (Fig.1J). Across all three subtypes, disruption of the axonal cytoskeleton emerged as a shared consequence of TDP-43 proteinopathy, whilst splicing defects were most prominent in MNs and V1 INs. The additional downregulation of chromatin and protein folding pathways specifically in MNs points to a broader and more severe transcriptional disruption in this subtype, consistent with its heightened vulnerability in ALS.

### Weighted gene co-expression network analysis reveals co-regulated transcriptional modules downstream of TDP-43 mislocalisation

To gain a systems-level understanding of the transcriptional changes observed across spinal neuron subtypes, we performed weighted gene co-expression network analysis (WGCNA) using the hdWGCNA package, which is specifically adapted for single-cell data (Morabito et al., 2023; Langfelder and Horvath, 2008). Day 22 and Day 40 annotated datasets were merged prior to network construction. Genes expressed in fewer than 5% of cells were excluded to reduce noise.

A key step in hdWGCNA is the construction of metacells, in which k-nearest neighbour (KNN) aggregation is used to identify groups of transcriptionally similar cells within the same condition and cell type, and their expression values are averaged to yield a denser metacell matrix. This aggregation step is necessary to reduce the inherent sparsity of single-cell count matrices, which would otherwise compromise network construction. The overall matrix density increased substantially from 0.05 in the raw count matrix to approximately 0.20 at k=25, with further incremental gains at higher values, confirming that metacell aggregation effectively recovered co-expression signal.

To select the optimal value of k, we systematically tested a range from 25 to 50 and evaluated each solution on four criteria: overall matrix density, cell type-specific matrix density, number of co-expression modules identified, and the size of the grey module i.e. the module to which genes are assigned when they cannot be meaningfully co-clustered. Cell type-specific density analysis showed that MNs and V1 INs consistently achieved the highest matrix densities across all k values, whilst V2 IN neurons showed lower densities throughout, likely reflecting their smaller cell numbers in the dataset. Across all metrics, k=35 provided a balance between sufficient matrix density and network resolution without over-aggregating cells, and was selected for all subsequent network construction and module identification.

Metacells were extracted specifically from the motor neuron cluster, pooling Day 22 and Day 40 cells, and a MN-specific co-expression network was constructed. This yielded 27 distinct gene modules containing a median of 180 genes, to which each gene in the network was unambiguously assigned (Fig.2A, B). Genes assigned to the grey module, representing genes that could not be robustly co-clustered with any module, were excluded from further analysis.

**Figure 2.**
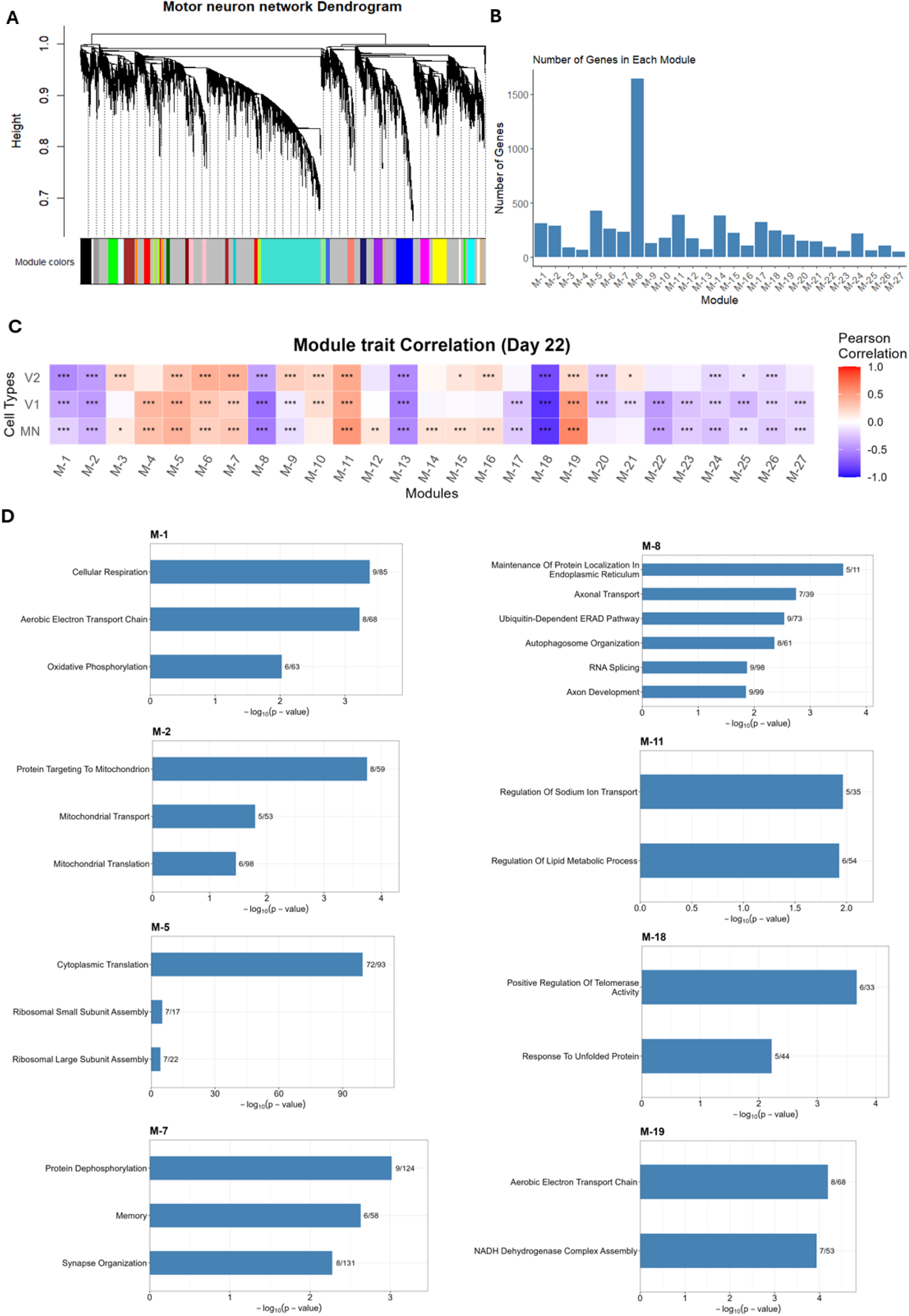
Weighted gene co-expression network analysis identifies TDP-43 mislocalisation-responsive modules in motor neurons. (A) Dendrogram showing hierarchical clustering of genes in the motor neuron co-expression network constructed using hdWGCNA. Each branch colour in the module colour bar represents a distinct co-expression module. A total of 28 modules were identified. Genes assigned to the grey module could not be unambiguously co-clustered and were excluded from downstream analysis. (B) Bar chart showing the number of genes assigned to each of the 27 co-expression modules. Module colours were replaced with numbers. The grey module has been removed. (C) Heatmap showing Pearson correlations between module eigengenes and TDP-43 condition (mislocalised = 1, nuclear = 0) across motor neurons (MN), V1 interneurons (V1), and V2 interneurons (V2). Red indicates a positive correlation (increased module activity in mislocalised cells); blue indicates a negative correlation (reduced activity in mislocalised cells). Asterisks denote statistical significance (* p < 0.01, ** p < 0.001, *** p < 0.0001). (D) Bar plots showing enriched GO terms in select modules. X-axis represents the −log10 adjusted p-value. The numbers on each bar plot indicate number of genes for a given GO term identified in the module / total number of genes in that GO term.

To identify modules whose activity is altered by TDP-43 mislocalisation, module eigengenes (MEs) were computed for each of the 27 modules. The ME summarises the overall expression state of a module as a single value per cell, analogous to the first principal component of module gene expression. Condition was binarised (mislocalised = 1, control = 0) and Pearson correlation was computed between each ME and the binarised condition variable for day 22 cells, with statistical significance assessed using a two-sided cor.test. Positive correlations identify modules with increased transcriptional activity in TDP-43 mislocalised MNs relative to controls, whilst negative correlations identify modules that are suppressed. We performed this analysis at day 22 to identify the earliest changes induced by TDP-43 mislocalisation (Fig.2C).

In MNs, modules 18, 8, and 13 were strongly negatively correlated with TDP-43 mislocalisation (r < −0.5), whilst modules 19 and 11 were strongly positively correlated (r > 0.5). Module 18 hub genes (highly connected genes in the module) included *STMN2*, *STMN4*, and *ELAVL3*, alongside genes encoding the nucleocytoplasmic transport (NCT) proteins RAN and RANBP1 and molecular chaperones DNAJB6 and DNAJC12, with GO enrichment pointing to unfolded protein response (Fig. 2D). NCT defects are documented in sporadic ALS, and the downregulation of RAN and RANBP1 here suggests that TDP-43 mislocalisation may compound existing NCT dysfunction (Kim et al., 2017). DNAJB6 is of particular interest, as it was recently shown to rescue FUS aggregation in a mouse model of ALS (Resnick et al., 2025); its downregulation in our model may therefore facilitate protein aggregate formation. Module 8 hub genes included *VCP*, an ALS-associated protein homeostasis factor, and *NEFL*, a structural component of MN axons and established ALS biomarker, with enrichment for proteostasis, axonal transport, RNA splicing, and autophagy terms (Fig. 2D). Module 13 lacked specific pathway enrichment but contained *XPO5*, an exportin required for microRNA biogenesis, as a hub gene.

Among upregulated modules, module 11 was enriched for sodium ion transport and lipid metabolism, an emerging theme in ALS biology, particularly in relation to cholesterol homeostasis (Hop et al., 2022, Agrawal et al., 2022) (Fig. 2D). Surprisingly, module 19 was enriched for aerobic electron transport chain, mitochondrial respiratory chain complexes I and IV, and NADH dehydrogenase activity (Fig. 2D), with hub genes comprising mitochondrially encoded transcripts including *NDUFB8.1, NDUFA4, COX7B*, and *COX7A2*. This observation likely reflects the elevated mitochondrial transcript levels observed in mislocalised MNs in our data.

V1 interneurons displayed a highly similar module correlation profile. Modules 18, 8, and 13 were again strongly negatively correlated (r < −0.5) and module 19 positively correlated (r > 0.5), with module 11 falling marginally below the correlation threshold (r = 0.49). Analysis of module 13 identified no enriched GO terms.

V2 interneurons showed downregulation of modules 18 and 1, the latter enriched for aerobic electron transport chain and oxidative phosphorylation terms (Fig. 2D), indicating mitochondrial stress. Notably, V2 INs did not show upregulation of module 19, suggesting an absence of the compensatory mitochondrial transcriptional response seen in MNs.

Across all three subtypes, modules 1 and 2 showed moderate negative correlation and was enriched for oxidative phosphorylation, mitochondrial translation and protein import terms, pointing to a sustained and broadly shared disruption of mitochondrial homeostasis downstream of TDP-43 proteinopathy.

Two further modules showed modest positive correlations across subtypes. Module 5 was upregulated in MNs and V1 INs, with weaker induction in V2 INs, and was enriched for cytosolic ribosomal protein subunits, both large and small, consistent with a UPR-driven compensatory increase in translational machinery, as discussed above (Pakos-Zebrucka et al., 2016). In V2 INs, module 7 showed a modest positive correlation and was enriched for synapse organisation and memory-associated terms, suggesting a subtype-specific transcriptional response that may reflect attempted synaptic compensation in the context of TDP-43 pathology.

### Correlating modules with TDP-43 mislocalisation

To determine which modules and pathways respond earliest to TDP-43 mislocalisation, we performed pseudotime trajectory analysis using *STMN2* cryptic exon (CE) inclusion as a proxy for the degree of TDP-43 nuclear depletion. *STMN2* CE inclusion is amongst the most sensitive molecular readouts of TDP-43 loss-of-function and is recoverable in single-cell protocols through its poly-A tail (Klim et al., 2019; Melamed et al., 2019). Each cell was assigned a mislocalisation score based on the ratio of CE-mapped reads to unspliced *STMN2* reads. Control nanobody-expressing cells registered near-zero scores, whilst NES nanobody-expressing cells displayed substantially higher scores with considerable within-condition variability (Fig.3A), capturing a spectrum of mislocalisation severity. Cells were then ordered along this pseudotime axis by cell type, and module eigengenes were plotted along the trajectory with Spearman rank correlations computed between eigengene and mislocalisation score.

**Figure 3.**
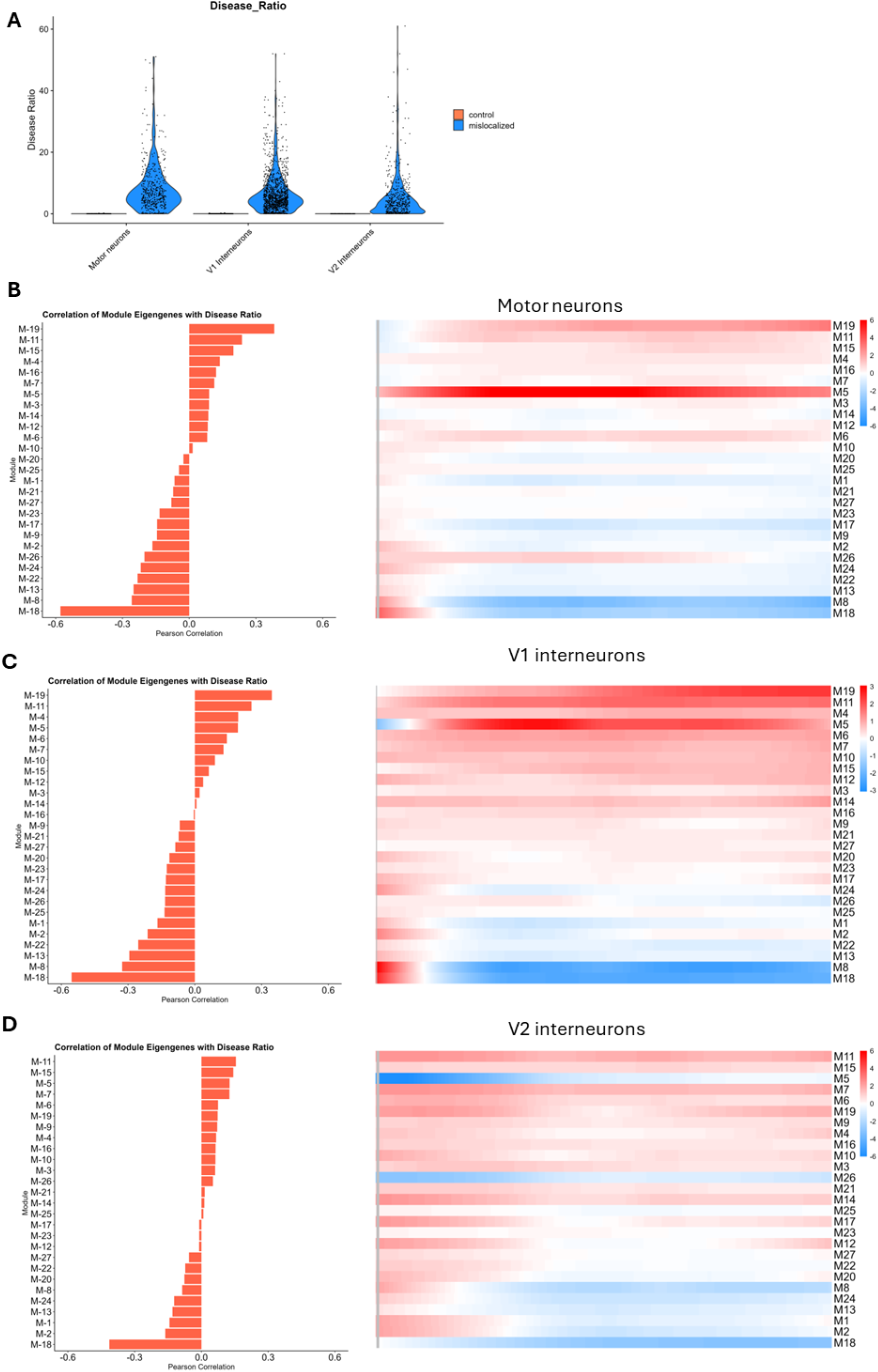
Pseudotime trajectory analysis orders cells along a TDP-43 mislocalisation gradient and identifies early-responding co-expression modules. (A) Violin plots showing the STMN2 cryptic exon disease ratio (cryptic exon read counts / total STMN2 read counts) per cell, plotted separately for MN, V1, and V2 interneurons. Control cells (orange) show near-zero disease ratios confirming effective suppression of cryptic exon inclusion under nuclear TDP-43 conditions. Mislocalised cells (blue) display a wide distribution of disease ratios, reflecting cell-to-cell variability in the degree of TDP-43 nuclear depletion. (B) Left panel: Bar charts showing Pearson correlations between module eigengenes and the STMN2 cryptic exon disease ratio pseudotime ranking, displayed for MNs. Left-side bars indicate negative correlations (modules downregulated with increasing mislocalisation); right-side bars indicate positive correlations. Right panel: Smoothed heatmaps showing module eigengene values plotted against cells ranked by increasing disease ratio (left to right) for MNs. Values were LOESS smoothed. Red indicates positive eigengene values and blue indicates negative values. The leftmost column represents the median eigengene value of each module in cells with nuclear TDP-43 (cells with Disease Ratio = 0), separated from the mislocalised cell values by a vertical grey bar. Modules that decline early along the pseudotime trajectory represent sensitive and primary responders to TDP-43 nuclear depletion, whilst those changing only at high disease ratios reflect late-stage transcriptional responses. (C,D) Similar plots to (B) for V1 and V2 interneurons respectively.

In MNs and V1 interneurons, modules 18 and 8 were significantly negatively correlated with pseudotime and their downregulation was detectable very early in the mislocalisation process (Fig.3B,C), identifying them as primary and sensitive responders to TDP-43 nuclear depletion. Module 19 was the most strongly positively correlated module overall, though it showed strong induction only at the furthest extreme of the mislocalisation score, suggesting that the compensatory mitochondrial transcriptional response is a late-stage event associated with severe TDP-43 pathology rather than an early adaptive response. V2 interneurons displayed attenuated dynamics of the module eigengene scores across the pseudotime (Fig.3D) in line with their more muted transcriptional response to TDP-43 pathology.

### Master regulator analysis identifies chromatin remodelling factors and RNA-binding proteins as key drivers of TDP-43-induced gene dysregulation

To identify the upstream transcriptional regulators driving the observed gene dysregulation, we performed master regulator analysis (MRA) on the MN co-expression network. The topological overlap matrix was reconstructed using a Signed Nowick approach to retain both positive and negative gene-gene correlations. Starting from 1,337 transcription factors and co-TFs annotated in the AnimalTFDB database (Hu et al., 2019), we first filtered for expression, retaining only those with normalised counts above 10 in at least 10 cells. For each TF, targets were defined by correlation strength: genes falling within the top or bottom 1% of all pairwise correlations in the network were assigned as positive or negative targets (regulons) respectively. TFs without any targets in either direction were excluded, and where the target set exceeded 1,000 genes, only the top 1,000 most positively or negatively correlated genes were retained. This yielded 799 TFs taken forward for master regulator analysis, with a median target set size of 171 genes (Fig. 4A).

**Figure 4.**
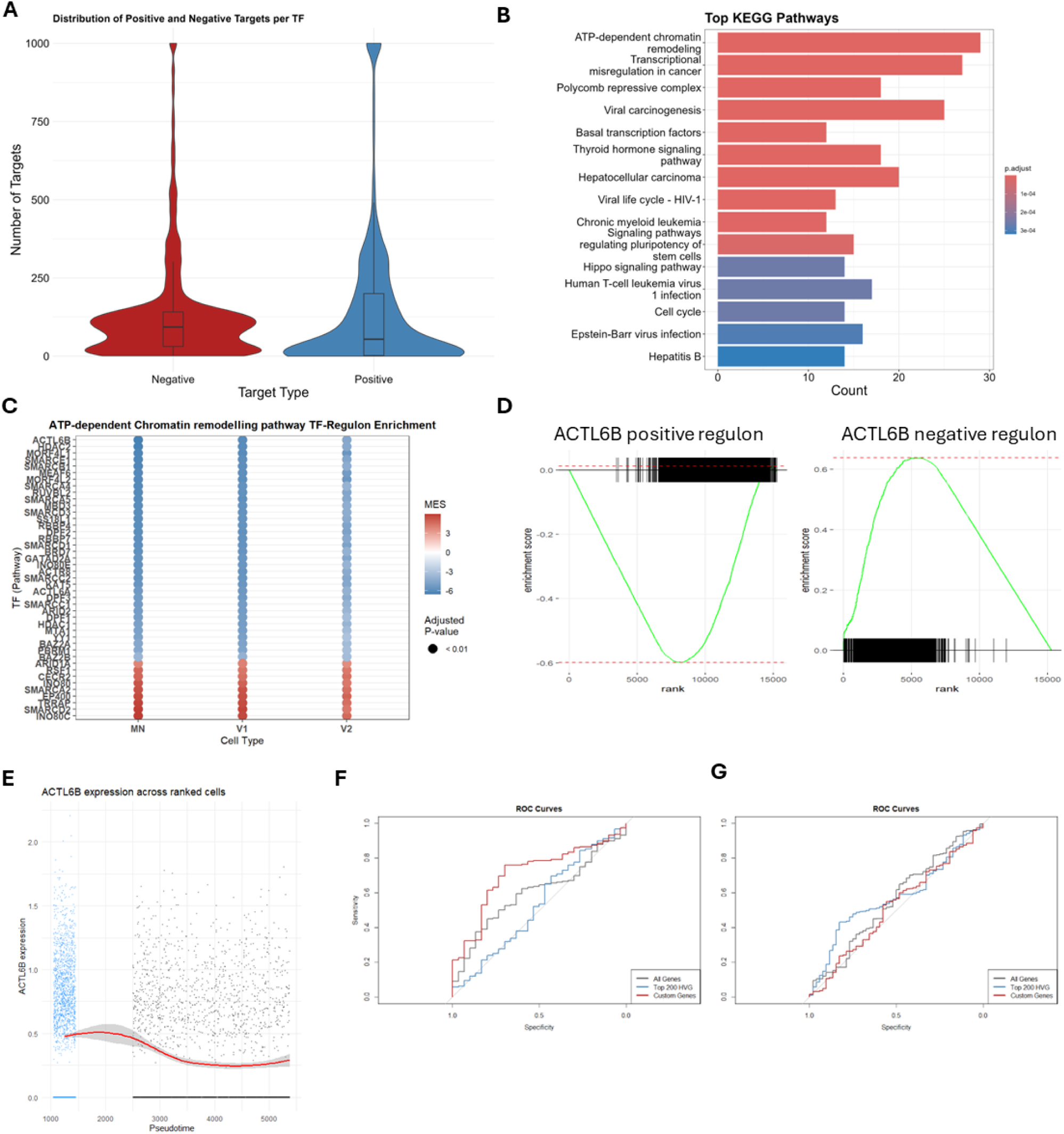
Master regulator analysis identifies ATP-dependent chromatin remodelling factors as drivers of TDP-43-induced gene dysregulation. (A) Violin plots showing the distribution of positive (blue) and negative (red) regulon sizes across the 799 transcription factors retained for master regulator analysis. Regulon size reflects the number of co-expression network genes positively or negatively correlated with each TF. (B) Bar chart showing the top KEGG pathways enriched amongst the 471 identified master regulators. ATP-dependent chromatin remodelling and polycomb repressive complex are the most significantly enriched pathways. Colour indicates adjusted p-value. (C) Dot plot showing the master enrichment score (MES; NES:positive regulon – NES:negative regulon) for each subunit of the ATP-dependent chromatin remodelling complex across MNs, V1, and V2 interneurons. Blue indicates reduced subunit activity in mislocalised cells; red indicates increased activity. Filled circles denote adjusted p-value below 0.01. ACTL6B shows the most consistently and strongly negative MES across all three cell types, identifying it as the most inhibited subunit of the complex following TDP-43 mislocalisation. (D) GSEA enrichment plots showing the enrichment of ACTL6B positive regulon genes (left) and negative regulon genes (right) across the ranked list of differentially expressed genes in mislocalised versus nuclear TDP-43 motor neurons. Positive regulon genes are enriched amongst downregulated genes and negative regulon genes amongst upregulated genes, confirming reduced ACTL6B transcriptional activity upon TDP-43 mislocalisation. (E) Scatter plot showing ACTL6B expression in individual cells ordered along the pseudotime axis, where pseudotime score is derived from the STMN2 cryptic exon disease ratio as a proxy for the degree of TDP-43 nuclear depletion. Blue dots represent cells with a disease ratio of zero, indicating retention of nuclear TDP-43. Grey dots represent cells with increasing disease ratios, reflecting a spectrum of TDP-43 mislocalisation severity. The red line denotes a smoothed expression trend (loess fit) with 95% confidence interval shown in grey shading. (F) ROC curves showing the performance of three gene sets: all genes, top 200 highly variable genes (HGV), and the custom ATP-dependent chromatin remodeller and polycomb repressive complex gene set (Custom Genes), in classifying ALS versus healthy samples in the NYGC spinal cord dataset. The custom chromatin gene set achieves the highest AUC (0.76), substantially outperforming the comparator gene sets. (G) ROC curves showing classification performance of the same three gene sets as applied to the NYGC cortical dataset. The chromatin gene set confers no improvement over comparator gene sets in cortical tissue, indicating that ATP-dependent chromatin remodelling dysfunction is a spinal cord-enriched feature of ALS pathology.

GSEA was then run on the ranked list of MN DEGs using each TF’s positive and negative target sets as gene sets, generating regulon enrichment scores in both directions. These were combined into a master enrichment score (MES; positive minus negative regulon score), with p-values integrated using Fisher’s method and adjusted by Benjamini-Hochberg correction. At an adjusted p-value threshold of 0.001, we identified 471 master regulators of TDP-43-induced gene dysregulation in MNs, of which 404 were inferred to be inhibited and 67 activated. In V1 INs, 404 TFs were inhibited and 74 activated; in V2 INs, 344 were inhibited and 40 activated, consistent with the more limited transcriptional response observed in this subtype.

GO enrichment analysis of the 471 MN master regulators identified significant enrichment of ATP-dependent chromatin remodelling and polycomb repressive complex components (Fig.4B). These same two groups were also enriched for the V1 and V2 master regulator TFs. Focussed analysis of the ATP-dependent chromatin remodelling group revealed multiple subunits of the neuronal BAF (nBAF) complex with reduced activity following TDP-43 mislocalisation in all three neuronal subtypes (Fig.4C). The top candidate within this group was ACTL6B (Fig.4C,D), also known as BAF53B, the neuron-specific actin-related subunit of the nBAF complex (Wu et al., 2007). ACTL6B expression was reduced at the earliest stages of the pseudotime trajectory, indicating that its downregulation is a primary rather than late-stage response to TDP-43 nuclear depletion (Fig. 4E). Strikingly, ACTL6B was identified as a master regulator across all three spinal neuron subtypes (Fig. 4C), suggesting that loss of nBAF complex activity is a broadly shared consequence of TDP-43 proteinopathy across the spinal cord.

### Chromatin remodelling complex downregulation is a feature of ALS patient spinal cord tissue

To determine whether the chromatin remodelling dysfunction identified in our model is relevant to human ALS, we examined ACTL6B expression in two publicly available ALS patient datasets. GSE76220 comprises RNA-seq data from laser-captured spinal MNs from ALS patients (Krach et al., 2018), whilst GSE126542 profiled cortical nuclei sorted based on TDP-43 expression status from ALS patient tissue (Liu et al., 2019). ACTL6B expression was downregulated in both datasets, providing independent validation that the master regulators identified by our MRA are dysregulated in human disease.

We then expanded this analysis to the full gene sets representing ATP-dependent chromatin remodellers and polycomb repressive complex components, using GSEA to assess coordinate downregulation. The spinal cord dataset GSE76220 showed significant downregulation of both gene sets (p < 0.01). However, no significant enrichment was observed in the cortical dataset GSE126542. One key difference between these datasets is tissue of origin: spinal cord versus cortex. This raised the possibility that chromatin remodelling dysfunction is more pronounced in spinal cord than in cortical tissue in ALS.

To test this directly, we used the NYGC ALS Consortium dataset, which contains transcriptomic data from both cortical and spinal cord tissue (Humphrey et al., 2023). We asked whether our chromatin gene sets could classify samples as ALS or healthy controls, benchmarking their performance against all genes and the top 200 highly variable genes. In the spinal cord dataset, the chromatin gene set significantly outperformed both comparators, achieving an area under curve (AUC) of 0.76 compared to 0.56 for the other gene sets (Fig.4F). However, no equivalent improvement was observed when the classification was applied to the cortical dataset (Fig.4G). These data suggest that transcriptional dysregulation of chromatin remodelling complexes is a specific and robust feature of ALS spinal cord pathology, and may reflect the selective vulnerability of spinal MNs to the epigenetic consequences of TDP-43 proteinopathy.

### ACTL6B knockdown impairs dendritic arborisation and disrupts chromatin regulatory programmes in motor neurons

To assess the functional consequences of ACTL6B loss in MNs, we delivered shRNAs via lentiviral transduction to MNs derive from healthy iPSCs, achieving greater than 75% knockdown of ACTL6B for both shRNAs tested (Fig.5A). At day 34 of differentiation, control MNs displayed complex axonal networks and highly arborised dendrites (Fig.5B). ACTL6B knockdown produced a visibly reduced dendritic complexity, with both shRNA-1 and shRNA-2 yielding statistically significant reductions in dendritic arborisation. Signs of cellular stress including axonal fragmentation and blebbing were also observed in the shRNA-1 condition (Fig.5B).

**Figure 5.**
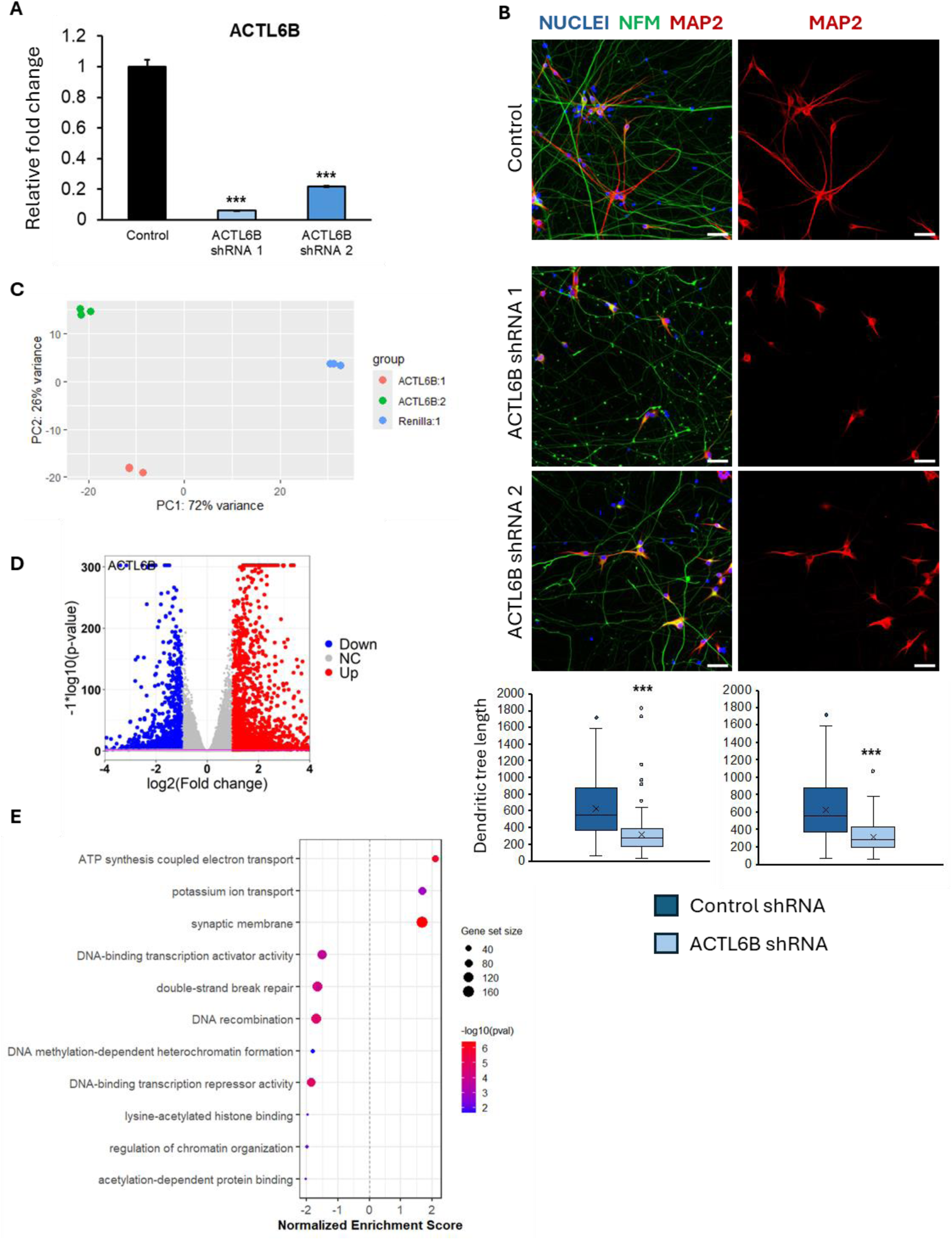
ACTL6B knockdown in post-mitotic iPSC-MN causes dendritic collapse. (A) ShRNA mediated knockdown of ACTL6B. ACTL6B expression was assessed using qPCR. HPRT1, GAPDH and RPL13 were used as housekeeping controls. Fold changes were normalised to control treatment. N=3 independent differentiations. N=3 independent differentiations. *** p < 0.001. P-values estimated using Student’s t.test. (B) Representative immunofluorescence images of iPSC-derived motor neurons at day 34 stained for nuclei (Hoechst, blue), neurofilament medium chain (NFM, green), and MAP2 (red). Top row shows control (Renilla shRNA) neurons with complex dendritic arbours. Middle and bottom rows show ACTL6B shRNA-1 and shRNA-2 conditions, respectively, demonstrating visibly reduced dendritic complexity. Scale bars indicate 50 um. Box plots show quantification of MAP2-positive dendritic tree length for shRNA-1 (upper) and shRNA-2 (lower) compared to Renilla control. N=3 independent differentiations. *** p < 0.001. P-values estimated using Student’s t.test. (C) PCA of bulk RNA-seq data from ACTL6B shRNA-1 (red), ACTL6B shRNA-2 (green), and Renilla control (blue) conditions. PC1 (72% variance) separates knockdown from control samples. Tight clustering of replicates within each condition confirms high reproducibility. ACTL6B shRNA-1 and shRNA-2 are separated along PC2 (26% variance), consistent with differences in knockdown efficiency between the two constructs. (D) Volcano plot showing differential gene expression following ACTL6B knockdown versus Renilla control. Downregulated genes are shown in blue and upregulated genes in red. ACTL6B is the most significantly downregulated gene, confirming effective knockdown. A broad transcriptional response is evident across both directions of change. (E) Dot plot showing GSEA normalised enrichment scores for selected pathways in ACTL6B knockdown motor neurons. Dot size represents gene set size and colour represents −log10 p-value.

To define the transcriptional consequences of ACTL6B depletion, we performed bulk RNA-seq 14 days after shRNA addition, using a non-targeting Renilla shRNA as control. PCA and hierarchical clustering confirmed clear transcriptomic separation between knockdown and control samples with high replicate concordance (Fig.5C). DESeq2 confirmed ACTL6B as one of the most significantly depleted transcript (Fig.5D). Expression of other nBAF complex members *CREST* (*SS18L1*) and *BRG1* (*SMARCA4*) was minimally affected, and no compensatory upregulation of *ACTL6A* was observed.

GSEA identified convergent pathway alterations consistent with ACTL6B’s role as the neuron-specific BAF complex subunit. Downregulated pathways were enriched for chromatin organisation, DNA demethylation, DNA-binding transcription repressor activity, and DNA repair, reinforcing that ACTL6B is required to maintain the chromatin regulatory programmes that sustain MN identity (Fig.5E). Upregulated pathways converged on mitochondrial respiration, possibly indicating a compensatory response (Fig.5E). Strikingly, inhibition of ACTL6B also upregulated potassium ion channels and synaptic membrane proteins, possibly reflecting aberrations in neuronal activity and communication (Fig.5E). Together, these data show that ACTL6B depletion suppresses chromatin regulatory and transcriptional programmes whilst driving a compensatory upregulation of mitochondrial metabolism, closely mirroring the transcriptional consequences of TDP-43 mislocalisation identified in our scRNA-seq analysis.

## Discussion

We used single-cell transcriptomics to characterise the transcriptional consequences of TDP-43 mislocalisation across spinal neuron subtypes in a physiologically faithful iPSC-derived MN model. Differential expression and pathway analysis revealed a conserved core response, downregulation of axonal cytoskeletal genes, RNA splicing factors, and chromatin regulatory programmes, across MNs, V1, and V2 interneurons, with MNs and V1 INs showing the broadest and most severe transcriptional disruption. Notably, the combinations of chromatin organisation and spliceosome terms were enriched specifically in MN downregulated pathways, pointing to a deeper transcriptional crisis in the cell type most vulnerable in ALS. Furthermore, hdWGCNA identified co-expression modules enriched for axonal cytoskeleton and mitochondrial respiration genes as early responders to TDP-43 mislocalisation, with their downregulation detectable at the earliest stages of the pseudotime trajectory. Master regulator analysis converged on ATP-dependent chromatin remodelling factors, with *ACTL6B* emerging as a top candidate across all three neuronal subtypes. Knockdown of *ACTL6B* in post-mitotic MNs produced dendritic simplification, axonal stress markers, and transcriptional suppression of chromatin and transcription regulatory pathways, closely phenocopying the consequences of TDP-43 mislocalisation.

Validation in public ALS patient datasets confirmed downregulation of ATP-dependent chromatin remodeller and polycomb repressive complex gene sets in laser-captured spinal MNs (GSE76220), whilst no equivalent enrichment was observed in sorted cortical nuclei (GSE126542). To test whether this difference reflects a genuine tissue-specific pattern, we used NYGC ALS Consortium data and asked whether our chromatin remodeller gene sets could classify ALS from healthy samples in spinal cord and cortex independently. The chromatin gene set substantially outperformed all-gene and highly variable gene comparators in the spinal cord, achieving an AUC of 0.76 versus 0.56, but conferred no classification benefit in cortical tissue. Supporting this, ACTL6B cryptic exon inclusion has been reported to be modest in FTLD-TDP post-mortem brain tissue compared to spinal cord, suggesting that TDP-43-dependent disruption of ACTL6B-mediated chromatin remodelling is more consequential in spinal cord than in cortex (Faura et al., 2025). Together, these data point to ATP-dependent chromatin remodelling dysfunction as a spinal cord-enriched feature of TDP-43 proteinopathy, consistent with the selective vulnerability of spinal MNs in ALS.

ACTL6B, also known as BAF53B, is the neuron-specific actin-like subunit of the nBAF chromatin remodelling complex, essential for ATP-dependent regulation of chromatin accessibility in post-mitotic neurons (Wu et al., 2007). During neuronal differentiation, ACTL6B replaces ACTL6A in the BAF complex, a switch mediated by miR-9* and miR-124 that marks the transition from progenitor to post-mitotic identity (Yoo et al., 2009). Assembled nBAF interacts with the calcium-dependent co-factor CREST to promote dendritic development and arborisation (Aizawa et al., 2004). Mis-splicing of ACTL6B is now well-established as a consequence of TDP-43 nuclear depletion, observed in iPSC-derived MNs and post-mortem ALS tissue across multiple independent studies (Irwin et al., 2024; Seddighi et al., 2024; Rothstein et al., 2025, Gansssauge et al., 2025). The cryptic exon inclusion in *ACTL6B* is in-frame and though the transcript appears resistant to nonsense-mediated decay, with cryptic peptides detectable at the protein level, a significant reduction in ACTL6B expression at both transcript and protein levels has been clearly demonstrated, indicating a net loss-of-function effect (Seddighi et al., 2024). Our scRNA-seq pseudotime analysis shows that *ACTL6B* downregulation occurs early in the mislocalisation process, identifying it as a primary rather than late-stage consequence of TDP-43 nuclear depletion.

Our *ACTL6B* knockdown data demonstrate that its loss in already-differentiated MNs is sufficient to reduce dendritic complexity and suppress chromatin regulatory programmes, without compensatory upregulation of ACTL6A or changes in other nBAF subunits CREST or BRG1. This is consistent with ACTL6B being required not only for dendritic development but for the ongoing maintenance of dendritic architecture in post-mitotic neurons, a distinction important in the ALS context, where depletion occurs in mature neurons rather than during development. Prior work showed that post-mitotic inhibition of *BRG1*, another nBAF subunit, similarly impairs dendrite morphology (Tibshirani et al., 2017), and that combined loss of ACTL6B, CREST, and BRG1 is observed in ALS post-mortem spinal cord tissue (Tibshirani et al., 2017). We have documented reduced dendritic complexity in TDP-43 mislocalised MNs (Ganssauge et al., 2025), and our data suggest that ACTL6B mis-splicing contributes substantially to this phenotype. Whether restoration of ACTL6B expression or blockade of its cryptic exon via antisense oligonucleotides can rescue dendritic defects in TDP-43 mislocalised MNs is an important question for future work.

A striking additional finding from our ACTL6B knockdown transcriptomics was the upregulation of potassium channel genes. Elevated potassium channel expression would be expected to increase potassium conductance across the neuronal membrane, shifting the resting membrane potential towards a more hyperpolarised state. This would raise the threshold for action potential firing and reduce neuronal excitability, producing a state of hypoactivity. Notably, hypoexcitability has been documented in degenerating motor neurons in ALS and has been proposed as a feature of late-stage disease that may paradoxically follow an earlier period of hyperexcitability (Devlin et al., 2015). The upregulation of potassium channels upon ACTL6B loss in post-mitotic MNs therefore provides a potential mechanistic link between nBAF chromatin remodelling dysfunction due to TDP-43 pathology and altered motor neuron electrophysiology in ALS MNs.

Neuronal BAF dysfunction in ALS is unlikely to be limited to ACTL6B. Rare loss-of-function mutations in *CREST* have been identified in sporadic and familial ALS patients (Chesi et al., 2013; Teyssou et al., 2014), and heterozygous CREST loss produces motor deficits, neuroinflammation, and neuromuscular junction denervation in mice (Cheng et al., 2019). Notably, nBAF suppression has also been observed in FUS-ALS murine neurons, in a context where cryptic exon inclusion is not the operative mechanism (Tibshirani et al., 2017), suggesting that multiple upstream insults converge on nBAF dysfunction in ALS. ACTL6B depletion via TDP-43-dependent mis-splicing may therefore represent one node in a broader pattern of nBAF complex disruption that drives transcriptional dysregulation and neurodegeneration across ALS subtypes.

Finally, it is notable that ACTL6B loss-of-function in humans and mice predominantly produces cognitive rather than motor phenotypes (Bell et al., 2019; Vogel-Ciernia et al., 2013). This likely reflects a difference between developmental and acquired loss of nBAF function. In ALS, ACTL6B depletion occurs in mature, fully differentiated MNs, a context in which the complex has distinct target genes and regulatory roles compared to the developing nervous system. Understanding the target gene repertoire of nBAF in adult human MNs will be essential for defining the full downstream consequences of ACTL6B loss in ALS.

## Acknowledgements

We would like to thank the University of Exeter Sequencing Center. This study was funded by an MNDA Pilot award, and an MRC Response Mode Grant to AB. This study was supported by the National Institute for Health and Care Research Exeter Biomedical Research Centre. The views expressed are those of the author(s) and not necessarily those of the NIHR or the Department of Health and Social Care.

